# Loss of Nucleotide Sugar Transporter (*AtNST*) gene function in the Golgi membranes impairs pollen development and embryo sac progression in *Arabidopsis thaliana*

**DOI:** 10.1101/2025.07.10.664101

**Authors:** Rimpy Diman, Pinninti Malathi, Ramamurthy Srinivasan, Shripad Ramchandra Bhat, Yelam Sreenivasulu

## Abstract

- <u>N</u>ucleotide <u>S</u>ugar <u>T</u>ransporters (NSTs) are transmembrane proteins which are localized in the Golgi membranes and transport nucleotide sugars from cytosol to Golgi lumen. Transported nucleotide-sugars serve as donors in post-translational modification of proteins/ lipids, and donate sugar moiety to growing carbohydrate chain on nascent protein/lipid molecule, a reaction catalyzed by the enzyme glycosyltransferase.
- Here we reported that a mutation in the *Arabidopsis thaliana* gene *At3g11320* coding for NST protein causes defects in male gametophyte development including collapsed, nonviable pollen. This mutation also caused impairment in the female gametophyte progression by arresting it at the functional megaspore (FM) stage.
- Further, the mutant phenotype including silique size and seed set was reverted when the cDNA of *AtNST* gene was over-expressed in the mutant back ground. No abortive ovules were found in the siliques from the complemented plants.
- The results suggest that *AtNST* (*At3g11320*) gene might be responsible for maintaining and regulating transport of nucleotide-sugars from cytoplasm to the Golgi lumen for the glycosylation of essential proteins/carbohydrates/lipids etc., which are necessary for both male and female gametophyte development stages. This study shed further light on the role of such nucleotide sugar transporters in plant reproductive development processes, at least in *Arabidopsis*.

## Introduction

Nucleotide sugar transporters (NSTs) transport nucleotide sugars from the cytosol to the Golgi/ER lumen (Gerardy-Schahn *et al.,* 2001, Handford *et al.,* 2006) where these nucleotide sugars act as donors of sugar residues in the glycosylation of proteins, lipids, and proteoglycans, as well as in the biosynthesis of plant cell wall polysaccharides. Glycosylation increases stability and solubility of proteins, assists protein folding and also helps in targeting proteins to their destinations. NSTs are antiport and type III transmembrane proteins (Eckhardt *et al.,*1999). NST proteins are 300-350 amino acids long, highly hydrophobic and are predicted to possess 6-10 trans-membrane α-helical spans for transferring substrates (nucleotide-sugars) across the membrane (Descoteaux *et al.,* 1995; Baldwin *et al.,* 2001; Norambuena *et al.,* 2002). In plants, NSTs were included as NST/triose phosphate translocator (TPT) superfamily proteins.

In *Arabidopsis*, the diploid microspore and megaspore mother cells undergo meiosis to produce a tetrad of micro- and mega-spores in the anthers and the ovules, respectively. The microspores, after releasing from the tetrad undergo an asymmetric mitosis to form a larger vegetative cell and a smaller generative cell. The generative cell again undergoes one more round of mitosis to produce two sperm cells (McCormick, 1993, 2004). This three-celled pollen grain is called the male gametophyte. On the female side, out of the four megaspores, only one survives as functional megaspore. This functional megaspore undergoes three rounds of mitotic divisions and subsequent cellularization to produce a seven-celled mature embryo sac, also termed the female gametophyte (Yang & Sundaresan, 2000; Drews and Yadegari, 2002). These gametophyte developmental processes are highly regulated and mediated by a number of genes. These genes act in highly regulated and coordinated manner, and mutations in gametophyte developmental genes severely affect gametogenesis. Although a large number of gametophyte mutants are available, the identity of genes and their mechanism of action in gametophyte and embryo development remain to be established

*Arabidopsis thaliana* genome contains more than 50 genes belonging to the NST gene family and is distributed in six clades (Knappe *et al.,* 2003; Bakker *et al.,* 2005). Out of these, only a few are functionally characterized. Most of the earlier characterized NST mutants in plants exhibited impaired plant growth and development. The rice Golgi NST mutant (*OsNST1*), *brittle culm14* (*bc14*), displays reduced mechanical strength due to decreased cellulose content and altered cell wall structure, and exhibits abnormalities in plant development (Zhanga *et al.,* 2011). So far, NST mutants have been mostly characterized as vegetative defect mutants whereas their role in reproductive development has not been much explored. Reyes *et al.,* (2010) reported *Atutr1* and *Atutr3* double mutant as male and female gametophyte progression defect mutant (Reyes *et al.,* 2010), whereas, individual mutants were not showing obvious phenotype.

In the present study, we report the results on characterization of a novel mutation in a nucleotide sugar transporter (*At3g11320*) gene (*AtNST*) from *Arabidopsis thaliana*. This NST protein was localized in Golgi membranes and transports nucleotide sugars from cytosol to its lumen. Mutation in this gene affects the transport of these sugars which leads to impair the glycosylation process and this impairment prevents the progression of the gametogenesis in male and female gametophytes which causes subsequent seed sterility.

## Materials and methods

### Plant Material and Growth Conditions

Development and screening of transferred DNA (T-DNA) insertion population of *A. thaliana* (ecotype Columbia) has been described earlier (Pratibha *et al.,* 2013). Briefly, the T-DNA promoter trap lines were generated by floral dip transformation (Clough and Bent, 1998) of *A. thaliana* with *Agrobacterium tumefaciens* strain GV3101 harboring a bidirectional promoter trap vector carrying a promoter-less GFP gene at the right border and a promoter-less GUS gene at the left border of T-DNA. The T_1_ seeds were screened for T-DNA insertion by selection on a medium containing kanamycin (50 mg l^−1^). The green seedlings were transferred to pots and allowed to grow under controlled conditions (20±1 °C, 60% relative humidity, 16 h light/8 h dark, under fluorescent illumination of 100 μmol m^−2^ s^−1^) and T_2_ seeds were collected. Each kanamycin resistant T_1_ plant was considered as an independent trap. T_2_ plants of each line were examined for the reporter gene expression or seed sterility. One of the lines showing seed sterility (designated as bitrap-636) was identified and analysed in this study.

### RNA isolation and RT-PCR analysis

Total RNA was isolated from different parts of plant using total RNA extraction kit (Real Genomics, Taiwan), treated with DNase (Invitrogen, USA) and quantified by nanodrop spectrophotometer, ND-1000 (Thermo Scientific, USA). First strand cDNA synthesis was done using Superscript III reverse transcriptase (Invitrogen, USA) with oligo dT primer. The constitutively expressed GAP C gene was used as the normalization control. Double stranded cDNA was amplified with gene specific primers NST-F (5’-TTTGCTT CGTTTCTTGGCTTCGCTGTA-3’) and NST-R (5’-ATGAAGATAGCGGC CAATGGCCGGTTTT-3’). PCR product was analyzed on 0.8% agarose gel.

### Phylogenetic analysis

For phylogenetic tree construction, the amino acid sequences of different nucleotide sugar transporter proteins for *Arabidopsis* homologous genes as well as the orthologous genes from different species were retrieved from NCBI (http://www.ncbi.nlm.nih.gov/protein/). The amino acid sequences were aligned in TM-coffee mode of PSI-coffee server (Chang et. al. 2012). The Neighbor-Joining tree was constructed using the bootstrap test (1000 replicates) with MEGA 6.0 (Tamura *et al.,* 2013).

### Identification of T-DNA flanking sequence

Isolation and identification of T-DNA flanking sequences was done through Genome walking approach as described earlier (Pratibha *et al.,* 2013; Sharma *et al.,* 2015). Briefly, Genome walker libraries were constructed by digesting DNA with *Dra*I, *EcoR*V, *Pvu*II or *Stu*I restriction enzyme followed by adapter ligation. PCR amplification was carried out according to the manufacturer’s instructions (Clontech, USA). Primary PCR product was diluted (1:250) and used as a template for nested PCR amplification using a nested adapter primer (AP2) and a nested gene-specific primer. Resolved nested PCR product was eluted using a GFX PCR DNA Gel Band Purification kit (GE HealthCare, UK), cloned in pGEM-T Easy vector (Promega, USA) and sequenced. The sequence retrieved after removal of the vector backbone was BLAST searched against the Arabidopsis genome database (TAIR, www.arabidopsis.org) to localize the T-DNA insertion site.

### Phenotypic characterization of pollen and ovules

The inflorescences were collected and fixed in ethanol-acetic acid (9:1) for 24 h and dehydrated in ethanol series (50, 70 and 90%) followed by dissection of ovary and anthers in the Hoyer’s solution (Anderson, 1954) under stereozoom microscope SMZ1500 (Nikon, USA) and mounted on a glass slide for 2 h under cover slip and were observed under DIC optics.

For 4, 6-diamino-2-phenylindole (DAPI) staining, pollen were isolated from mature flowers and placed in 400 μl DAPI staining solution (4’, 6-diamidino-2-phenylindole in sodium phosphate (0.1 M; pH 7), 1 mM EDTA, 0.1% triton X-100, 0.4 mg/ml DAPI; high grade, Sigma) in micro centrifuge tube. After brief vortexing and centrifugation, pollen pellet was transferred to a microscope slide and viewed by light and by UV light under fluorescent microscope and images were captured.

For pollen viability analysis, pollen was collected from mature flowers and fixed in ethanol: glacial acetic acid (3:1) for 2 h. Stamens were excised and stained with a modified Alexander’s stain as per Peterson *et al.,* (2010). Nonviable pollens were distinguished from viable pollens by their purple colour observed under fluorescent microscope and images were captured with an Axioplan CCD camera using Axiovision software (AXIO imager M1, Carl Zeiss, GmbH, Germany).

For histological studies, the inflorescence was fixed in FAA (formaldehyde:acetic acid: 50% ethanol::1:1:18) for three days, dehydrated in t-butyl alcohol series, infiltrated and embedded in paraffin wax (m.p. 56 to 58°C). Sections of 12 µm thick were cut using rotary microtome (Shandon Finsse ME, Thermo Electron Corp., UK), stained with 0.5% toludene blue and mounted in DPX (Distrene, 8 to 10 g ;British resin product). Prepared sections were observed under fluorescent microscope under DIC optics.

For the scanning electron microscopy (SEM) analysis, mature pollen were mounted on aluminum stubs, carbon/gold coated using E-1010 sputter coating unit and observed under Scanning Electron Microscope (S-3400N Hitachi, Japan). The images were captured digitally.

### Complementation analysis

Full length cDNA of *At3g11320* gene (927 bp) was amplified by using NSTpcamF (5’-CCAGATCTGATGAAGATAGCGGCCAATGGCCG-3’) and NSTpcamR (5’-CCACTAGTTTTGCTTCGTTTCTTGGCTTCG-3’) primers from the total RNA of wild type Arabidopsis inflorescence converted into cDNA by using Superscript reverse transcriptase (Invitrogen,. USA). Resultant amplified fragment cloned in pGEMT-Easy vector (Promega, USA) and again sub-cloned into binary vector pCAMBIA1302. Resultant construct was mobilized into homozygous *Atnst* mutant plants by *Agrobacterium*-mediated infiltration (Bechtold & Pelletier, 1998). Transformants were selected on MSA plates supplemented with hygromycin (20 mg/ml) and genotyped by PCR using NSTpcamF and NSTpcamR primers and phenotype also analysed for its complementation.

### Cellular localisation of *AtNST* gene expression

Binary expression vector (pCAMBIA1302: *AtNST*) construct was mobilized into *Agrobacterium* GV3101 using the standard transformation techniques (Sambrook *et al.,* 1989). Fresh onion epidermal peels were re-suspended in *Agrobacterium* solution consisting of 5% (g/v) sucrose, 100 mg acetosyringone and 0.02% (v/v) Silwet-77 for 6–12 h at 28°C. After that the onion peelings were shifted to a Petri dish containing MS agar plates and incubated for 1–2 d. After washing with distilled water, onion epidermal cell layers were placed on glass slide and screened for the GFP expression localisation using laser scanning confocal microscope (Zeiss, LSM510 META, GmbH, Germany) equipped with a Zeiss Axiovert 200 M inverted microscope.

## Results

### Isolation, identification and characterization of *AtNST* gene mutant

Screening of in-house developed T-DNA insertion mutant lines of *A. thaliana* identified a number of seed sterile mutants. Among these, a mutant line with short siliques, abortive ovules and high seed sterility (82%) was observed and named as Bitrap-636 (Fig. 1). Aborted ovules appeared like white spots in between well developed seeds in mature siliques (Fig. 1F, G). In order to identify which of two gametes contributes towards seed sterility, reciprocal crosses were made between mutant and wild type plants and the siliques were examined for seed sterility. A seed sterility of 47.21% was observed when flowers of the mutant plants were pollinated with the wild type pollen, whereas, the reciprocal cross showed 55.64% seed sterility (Table 1). These results confirmed that the mutation leads to defects in both female and male gametes. To identify the nature of defect that leads to seed sterility, developmental pattern of both male and female gametophytes was examined in the Bitrap-636 mutant. Microscopic analysis of anthers revealed 84% of pollen (out of 347 pollen) was either collapsed or deformed and 82% ovules (out of 929 ovules) were abortive in the Bitrap-636 mutant and the rest were normal (Table 2).

**Figure 1:**
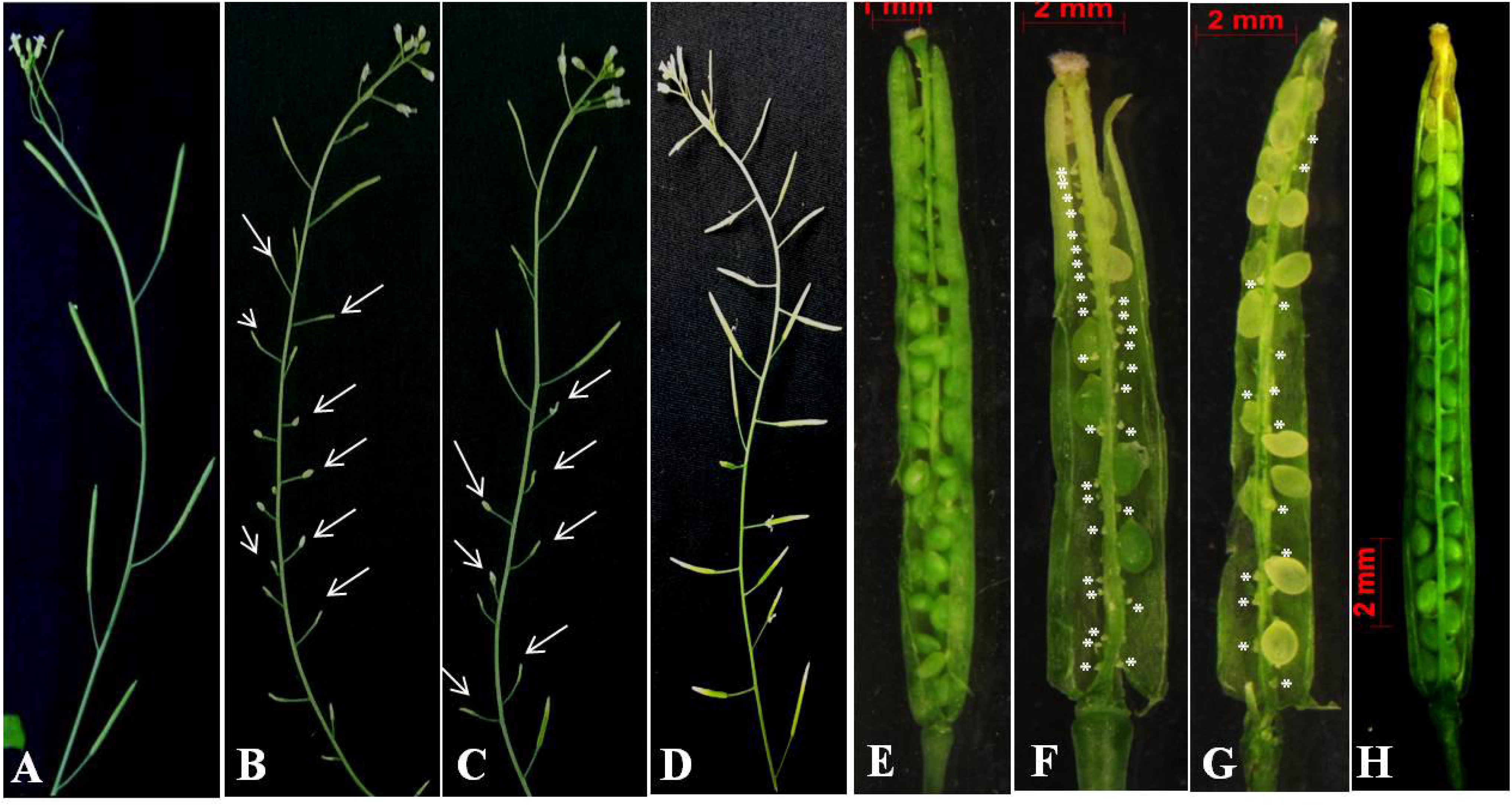
Phenotype of *Atnst1-1* and *Atnst1-2* mutant and complemented plants (A-D). (E-H) Silique showing seed sterility in WT and mutant lines, (A) wild-type, (B) *Atnst1-1* mutant, (C) *Atnst1-2* mutant. Mutants show fewer and shorter silique as compared with wild-type. (E) Silique from WT showing full seed set, (F) and (G) siliques from *Atnst1-1* and *Atnst1-2* mutants, respectively, showing some of the ovules developed into seeds and the others aborted (highlighted with asterisks). (D) and (H) Inflorescence and silique from the mutant plant complemented with the cDNA of *AtNST* gene, showing complete complementation

**Table 1.** Seed sterility in reciprocal crosses between wild type and *Atnst* mutant *Arabidopsis* plants.

| **Genotype** | **Number of** | **Aborted ovules** |
| --- | --- | --- |
| **Female X male** | **ovules observed** | **(%)** |
| **WT (self)** | 290 | 3 |
| **WT X *Atnst*/+** | 273 | 55.64 |
| ***Atnst*/+ X WT** | 315 | 47.21 |
| ***Atnst*/+ X *Atnst*/+** | 405 | 67.15 |

**Table 2.** Details of pollen and ovule abortion in *Atnst* mutant.

| **Genotype** | **Pollen** |  |  | **Ovules** |  |  |
| --- | --- | --- | --- | --- | --- | --- |
|  | **Total**<br>**number**<br>**checked** | **Viable**<br>**(%)** | **Non- viable**<br>**(%)** | **Total**<br>**number**<br>**checked** | **Normal**<br>**(%)** | **Aborted**<br>**(%)** |
| **WT** | 523 | 94.45 | 5.55 | 1288 | 96.62 | 3.38 |
| *Atnst1-1*<br>(homozygous) | 347 | 16 | 84 | 929 | 18 | 82 |

Genome walking approach was used to identify the T-DNA flanking sequences in the Bitrap-636 mutant. Primary and subsequent nested PCR with left border (LB) specific primers (TL GUS and TL GUSN) and adapter specific primers (AP1 and AP2) gave 900 bp and 700 bp amplified products (Supplementary Fig. S1), which were cloned and sequenced. BLAST search of the 800 bp sequence against TAIR database showed perfect match with *At3g13730* gene region and indicated that the T-DNA is inserted in the second intron of *At3g13730* gene (*AtCYP90D1*). To further confirm the T-DNA location, primers were designed from the T-DNA region and from either side of the putative insertion site (Supplementary Fig. S2A). PCR with primer pairs BLB - CYP90DR Ins-f gave amplicons of expected size (1.6 kb and 500 bp). However, the primer pair BRB - Ins-r gave longer than the expected amplicon; instead 300 bp an amplicon of 1.8 kb was obtained (Supplementary Fig. S2B). Cloning, sequencing and subsequent BLAST analysis of 1.8 kb fragment against TAIR database revealed 100% match with the nucleotide sugar transporter gene (*At3g11320*) region corresponding to 106 bp of 7^th^ exon of the gene, 371 bp of 3’UTR and 731 bp of intergenic region. These results suggested that certain rearrangement has been taken place during the T-DNA insertion. The T-DNA initially inserted in reverse orientation (left boarder in sense orientation with reference to the nucleotide sugar transporter gene (*At3g11320*). From there the T-DNA along with 1208 bp (106 bp of 7^th^ exon, 371bp of 3’UTR and 731 bp of intergenic region) inversely translocated into the 2^nd^ intron of *CYP90D1* gene (*At3g13730*) on the same chromosome. Mutation in *AtNST* was further confirmed through PCR using primer pair P1 and P2 (Fig. 2.1). Amplification of the expected size fragments i.e. 1.67 kbp and 464 kbp, respectively, confirmed the mutation in *At3g11320*. The T2 generation of *AtNST* mutant plant were analyzed by PCR to identify and isolate homozygous mutant plants. PCR with the primer pair P1 and P2 gave 1.67 kbp amplicon only in some of the progenies, whereas, in others an additional 464 kbp amplicon was also amplified (Fig. 2.1). In some of the progenies both the amplicons were amplified. Amplification of both the fragments confirmed their heterozygosity for the deletion. Amplification of 464 kbp only confirmed their homozygosity for the deletion mutation and amplification of 1.67 kbp amplicon represents WT.

**Figure 2:**
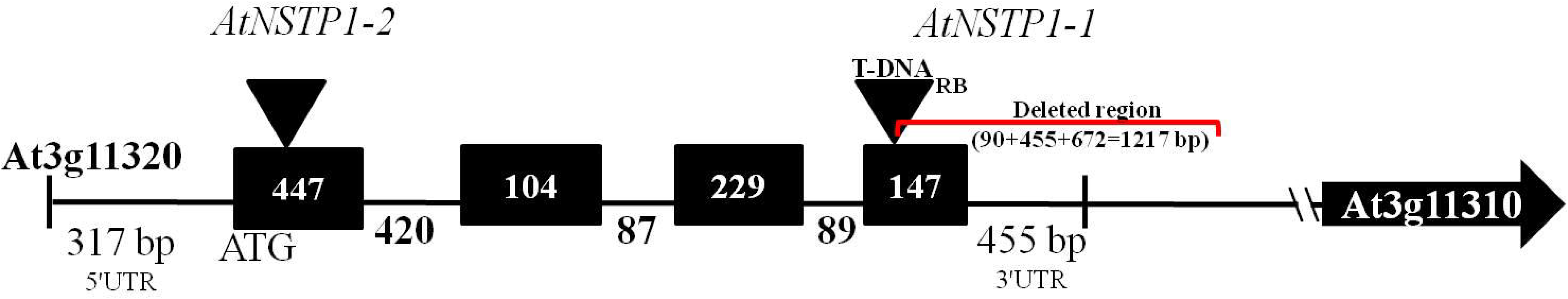
Diagrammatic representation of T-DNA insertion followed by translocation to a new location leading to deletion in the *AtNST* (*At3g11320*) gene.

Possible rearrangements are depicted in Supplementary Fig. S3. Due to this rearrangement two genes *AtCYP90D1* (*At3g13730*) and *AtNST* (*At3g11320*) were disturbed in the Bitrap-636 mutant line. Either of the gene mutation or both may be responsible for the mutant phenotype. In order to resolve this individual SALK lines having T-DNA insertion in each of these genes i.e. *AtCYP90D1* (SALK_042932) and *AtNST* genes (CS_358365) were obtained from ABRC and analyzed for the phenotypes (Supplementary Fig. S4A). In these SALK lines, T-DNA insertions were confirmed by PCR amplifications by using the gene specific primers. Two fragments of 1kb and 3 kb were amplified in SALK_042932, which showed that T-DNA is in hemizygous condition and confirmed the T-DNA insertion in the 1^st^ exon of the CYP90D1 gene (Supplementary Fig. S4B). Similarly amplicons of 1.5 kb, 2.8 kb amplified in CS_358365 showed that the T-DNA is in hemizygous condition and confirmed T-DNA insertion in the 1^st^ exon of the NST gene (Fig. 2). Here onwards, Bitrap-636 mutant having T-DNA insertion in *NST* gene and the NST gene SALK line (CS_358365) are referred as *AtNST1-1* and *AtNST1-*2, respectively. Silique phenotypes and seed fertility were analyzed in these SALK lines. Fully filled siliques were found in SALK_042932 (*CYP90D1*), whereas, CS_358365 (*NST*) line showed significant seed sterility with reduced seed set. Further, deformed, collapsed and aborted pollen were also found in anthers as in the mutant (Bitrap - 636). These results reconfirmed that mutation in the nucleotide sugar transporter gene is responsible for the mutant phenotype in Bitrap-636 mutant.

### *At*NST protein is a part of the nucleotide sugar transporter protein family

Gene coding for nucleotide sugar transporter protein occur as multi gene family with more than 40 members in *A. thaliana*. The *NST* gene *At3g11320* is one of the 40 sugar transporter genes and has four exons and three introns. It is predicted to encode a protein of 308 amino acids with a molecular weight of 33.8 kDa (iso-electric point of 10.2). A phylogenetic tree was constructed with *Arabidopsis AtNST* genes and rice, *C. elegans*, human and yeast *NST* genes (Fig. 3). In this four of *Arabidopsis* nucleotide sugar transporters i.e. *At5g05820*, *At3g10290*, *At5g04160* and *At1g12500* fall in a clade belonging to *At3g11320* gene group along with the two rice orthologs i.e. *Os03g0286300* and *Os05g0121900*. Homology search analysis at amino acid level between the members of this clade is presented in Table 3. *At3g11320* shares high homology (95%) with *At5g05820* and the least homology (65%) with *At1g12500.* A NCBI BLAST search of this protein revealed that it has a significant homology (85%) with *Oryza sativa* (*Os03g0286300*) gene which also encodes a 322 amino acid nucleotide sugar transporter protein with a number of trans-membrane domains (TMD) (Ohyanagi *et al*., 2006). *At3g11320* also contains about 10 transmembrane, inner- and outer-membrane domains (Fig. 4). Generally, these TMDs form a channel across the membrane for transport of nucleotide sugars from cytosol to Golgi lumen. Analysis of protein sequence with Euk-mPLoc 2.0 programme revealed that this protein has chloroplast and Golgi membrane localization signals and might serve to transport nucleotide sugars from cytosol to these organells to supply sugars for glycosylation.

**Figure 3:**
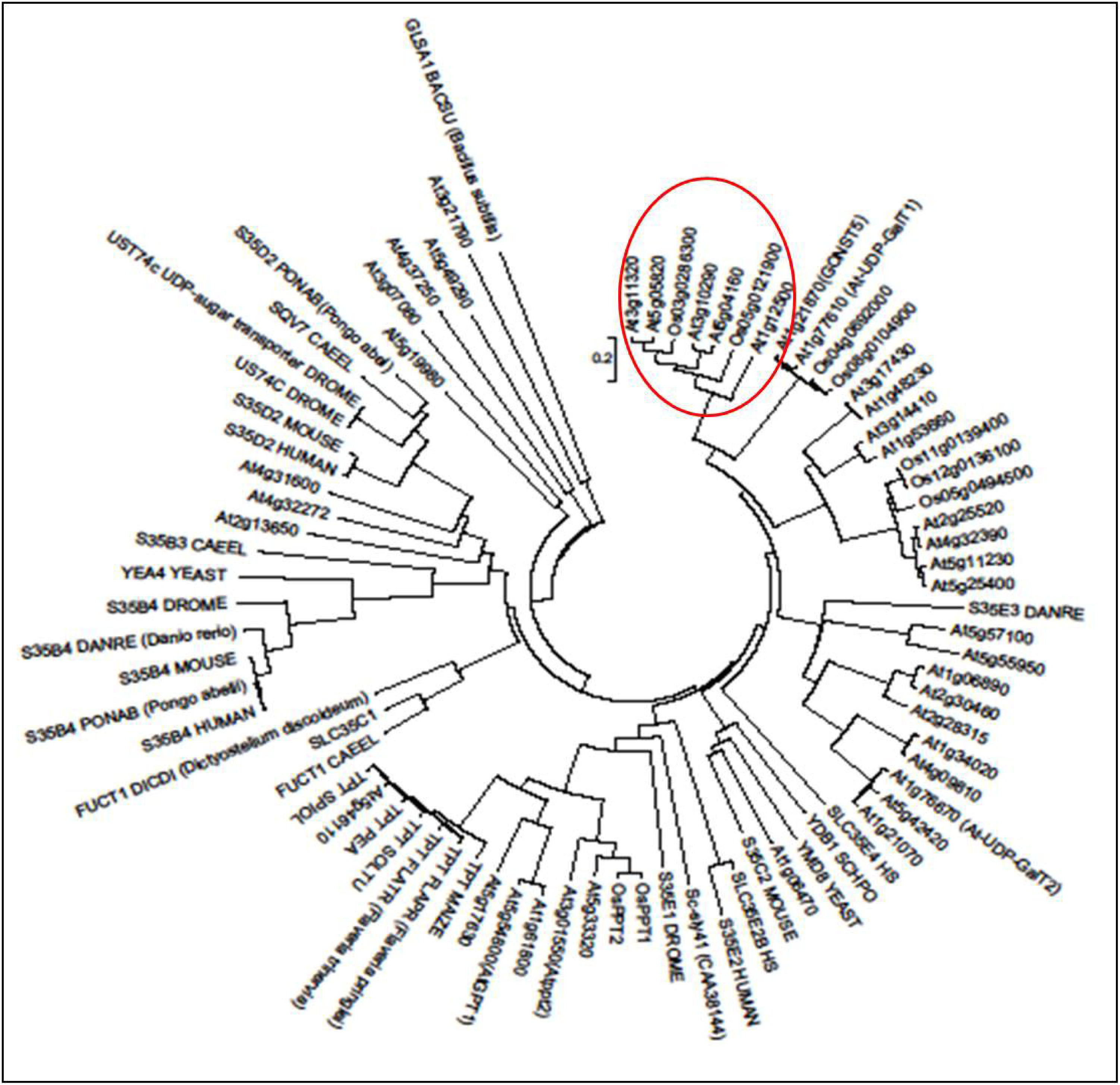
Phylogenetic analysis of the nucleotide sugar transporter proteins of *Arabidopsis thaliana* and *oryza sataiva*.

**Figure 4:**
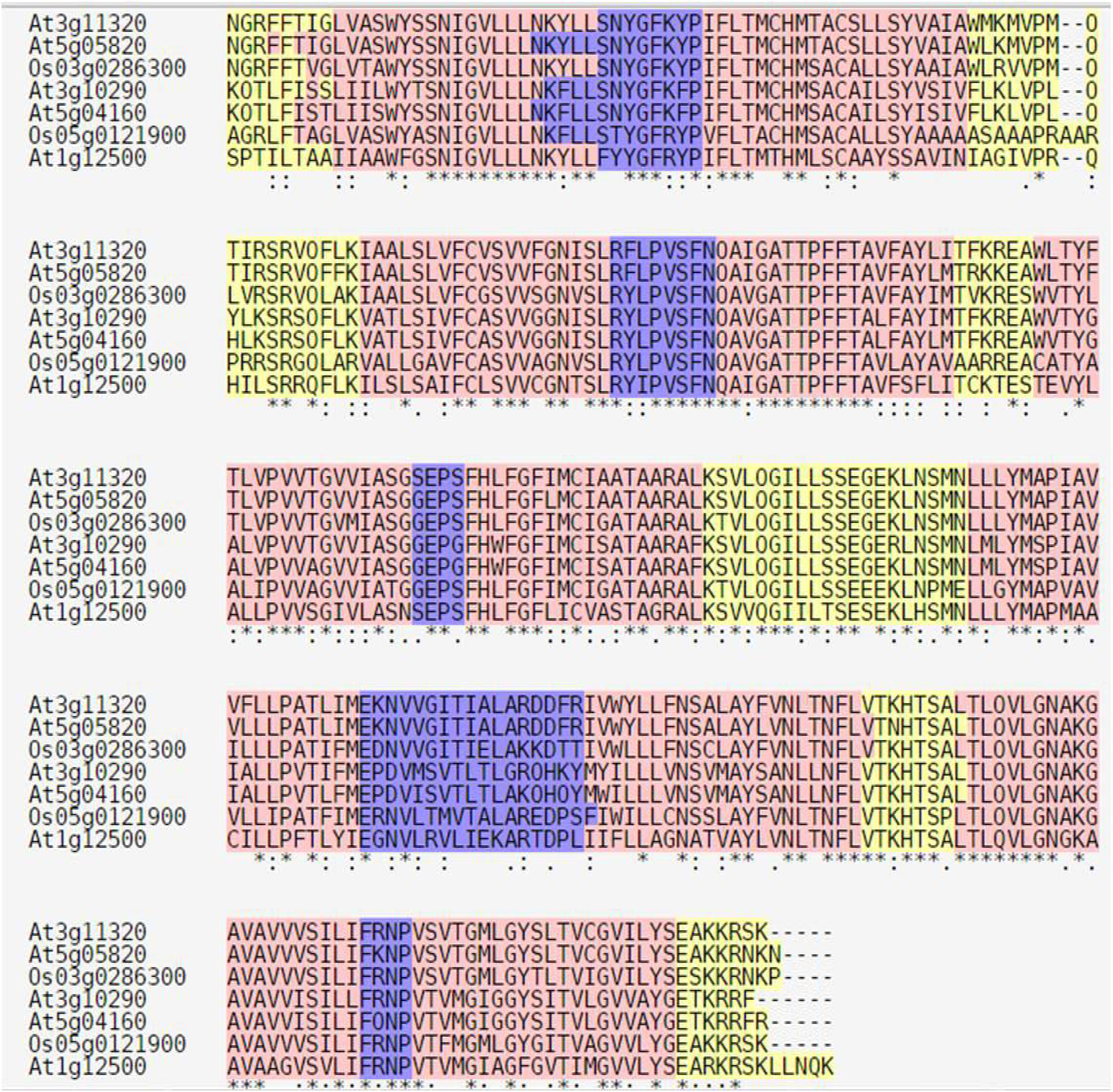
Multiple alignment of different NST proteins of *Arabidopsis thaliana* and *Oryza sativa.* Transmembrane domains are highlighted in pink and highly similar regions are highlighted in violet colour.

**Table 3.**
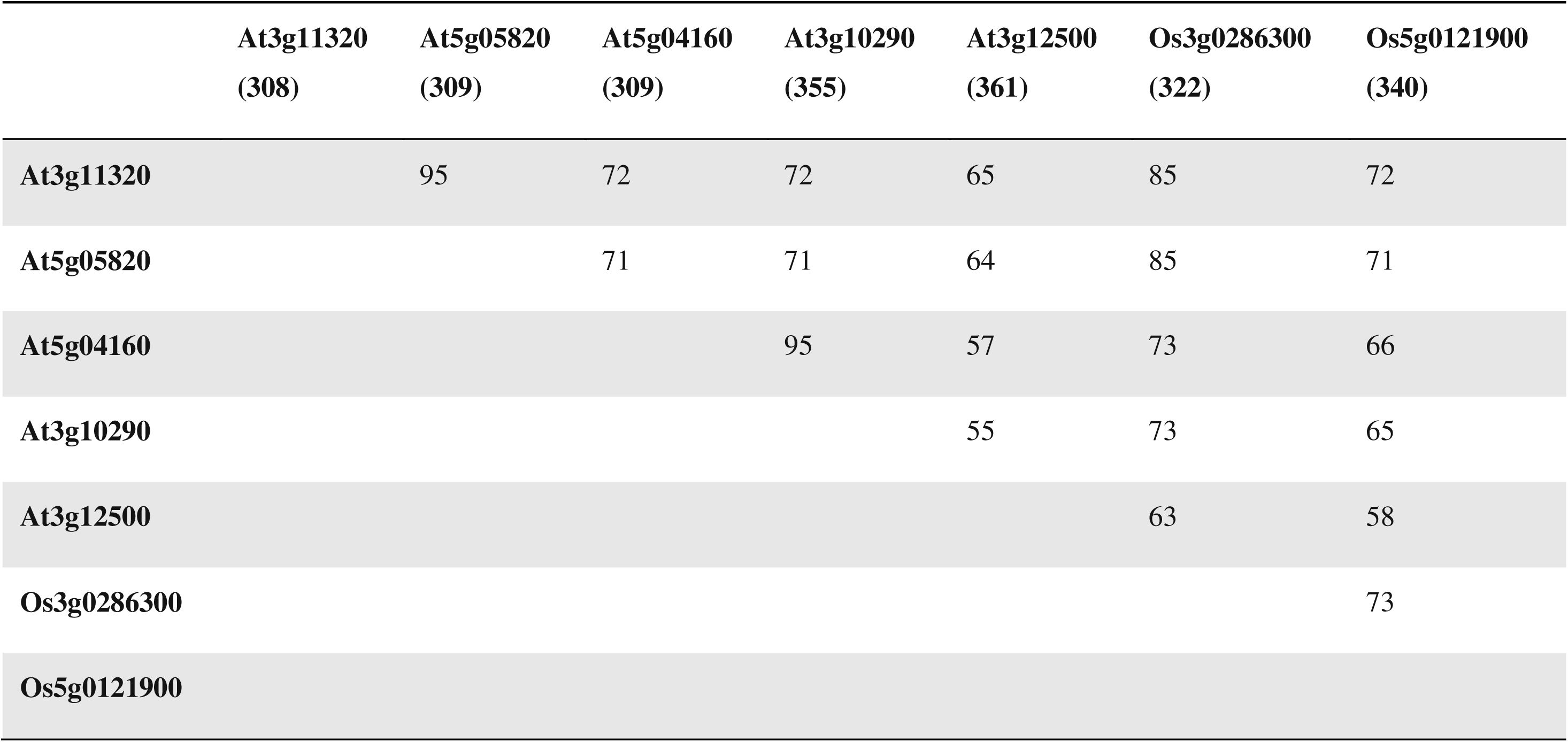
Homology of nucleotide sugar transporter proteins of Arabidopsis and rice. The values indicate the percentage amino acid identity between protein sequences. The length of putative polypeptide (number of amino acid residues) is given in parentheses.

|  | **At3g11320**<br>**(308)** | **At5g05820**<br>**(309)** | **At5g04160**<br>**(309)** | **At3g10290**<br>**(355)** | **At3g12500**<br>**(361)** | **Os3g0286300**<br>**(322)** | **Os5g0121900**<br>**(340)** |
| --- | --- | --- | --- | --- | --- | --- | --- |
| **At3g11320** |  | 95 | 72 | 72 | 65 | 85 | 72 |
| **At5g05820** |  |  | 71 | 71 | 64 | 85 | 71 |
| **At5g04160** |  |  |  | 95 | 57 | 73 | 66 |
| **At3g10290** |  |  |  |  | 55 | 73 | 65 |
| **At3g12500** |  |  |  |  |  | 63 | 58 |
| **Os3g0286300** |  |  |  |  |  |  | 73 |
| **Os5g0121900** |  |  |  |  |  |  |  |

### Expression pattern of *At3g11320* (*AtNST*) gene in *Arabidopsis*

According to *Arabidopsis* eFP browser and GENEVESTIGATOR analysis, this gene is expressed in stem, leaves, inflorescences and siliques and a high level expression is reported in roots of seedlings, developing anthers and in mature pollen (Supplementary Fig. S5). Expression of *At3g11320* gene was tested in wild type and mutant (hemizygous and homozygous) lines through RT-PCR. In wild type, *At3g11320* transcripts were found in silique, inflorescence, stem and leaf whereas hemizygous mutant lines showed reduced levels of transcripts, but no transcripts were detected in homozygous mutants (Fig. 5).

**Figure 5:**
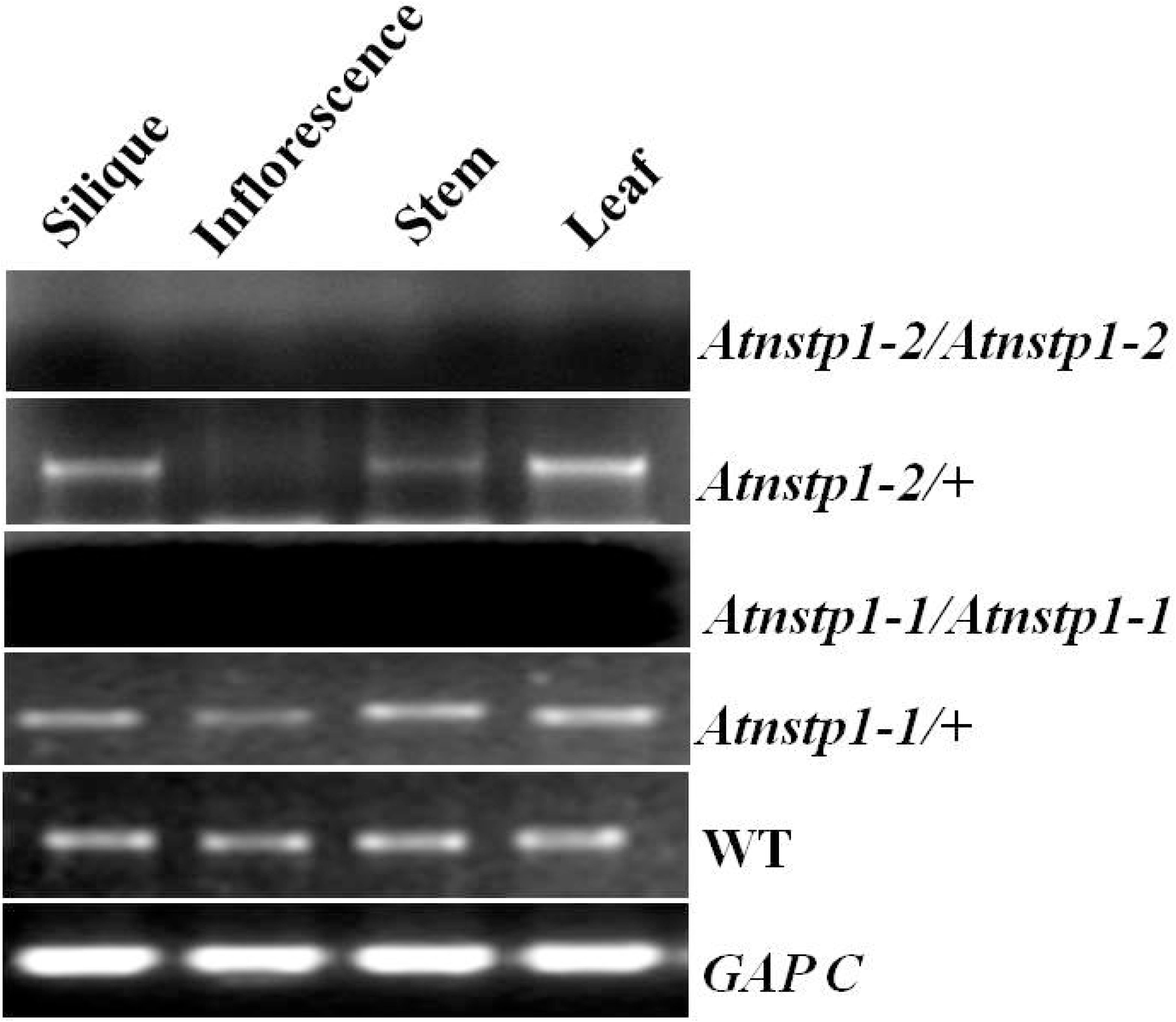
RT-PCR analysis of expression pattern of *At3g11320* gene in *Atnst1* mutants and wild type Arabidopsis.

### At3g11320 protein is localized in the Golgi membranes

In order to investigate the sub-cellular localization of At3g11320 protein, a construct was prepared in pCAMBIA1302 vector as a translational fusion with GFP and transiently expressed in the epidermal cells of the onion peelings. Laser Scanning Confocal microscopy analysis of these peelings confirmed the localization of GFP expression in the intracellular organs possibly in the ER and Golgi (Supplementary Fig. S6). *In silico* analysis of domain search by using Euk-m PLoc 2.0 programme (Chou *et al.,* 2010) was also localised this nucleotide sugar transporter protein in the membranes of endoplasmic reticulum. Conserved domain search by using NCBI showed for the presence of MKIAANGRFF, a domain related to trios-phosphate transporter (TPT) family protein. Parsons *et al.,* (2012) identified the protein encoded by *At3g11320* gene in the Golgi proteome in *Arabidopsis* which further validated its localization in Golgi membranes. All these evidences confirmed that *At3g11320* encodes a nucleotide sugar transporter which belongs to triose phosphate transporter family; this family includes transporters with specificity for triose phosphates.

### Male and female gametophyte development is affected in *Atnst* mutant

As crossing results revealed that male gametophyte is more affected and contributed more to seed sterility in *Atnst* mutant, detailed morphological analysis of pollen development was performed in the selected mutant. Tissue cleared anthers analysed with differential interference contrast (DIC) microscopy showed that a large proportion of pollen grains was deformed and collapsed (Fig. 6B and 6C). Upon Alexander’s staining, collapsed pollen grains (80%) stained purple and were therefore inferred as nonviable (Fig. 7 and Table 4). In contrast, the larger and normal-looking pollen (about 20% of the pollen) stained red similar to wild type pollen and were counted as viable (Fig. 7B and 7C). In order to assess cell cycle progression, number of nuclei was assessed in *Atnst* mutant pollen by staining with DAPI. Wild type pollen had three clear DAP-stained nuclei (Fig. 8A) whereas collapsed and degenerated pollen from the mutant plant did not have clear nucleus and showed diffused DAPI staining (Fig. 8B-F). Scanning electron microscopy analysis of the mutant pollen was performed to identify the structure of the sporopollenin/exine wall. All the pollen grains including normal as well as collapsed had normal reticulate pattern of outer exine wall (Fig. 8G-I). Histological analysis of different stages of mutant and wild type anther development using toluidine-blue (0.05%) staining was performed to investigate the exact developmental stage at which the phenotypic defects manifest (Fig. 9). The differences between *Atnst* mutant and wild type appear from stage 8 after release of microspores from the tetrads (Fig. 9). Up to the tetrad stage anther development was similar in both wild type and mutant. However, at stage 8, more than 80% microspores appeared mis-shaped, shrivelled, collapsed and defective in the mutant in comparison to wild type in which they were spherical and fully cytoplasmic. Only a small fraction of pollen in the mutant appeared normal.

**Figure 6:**
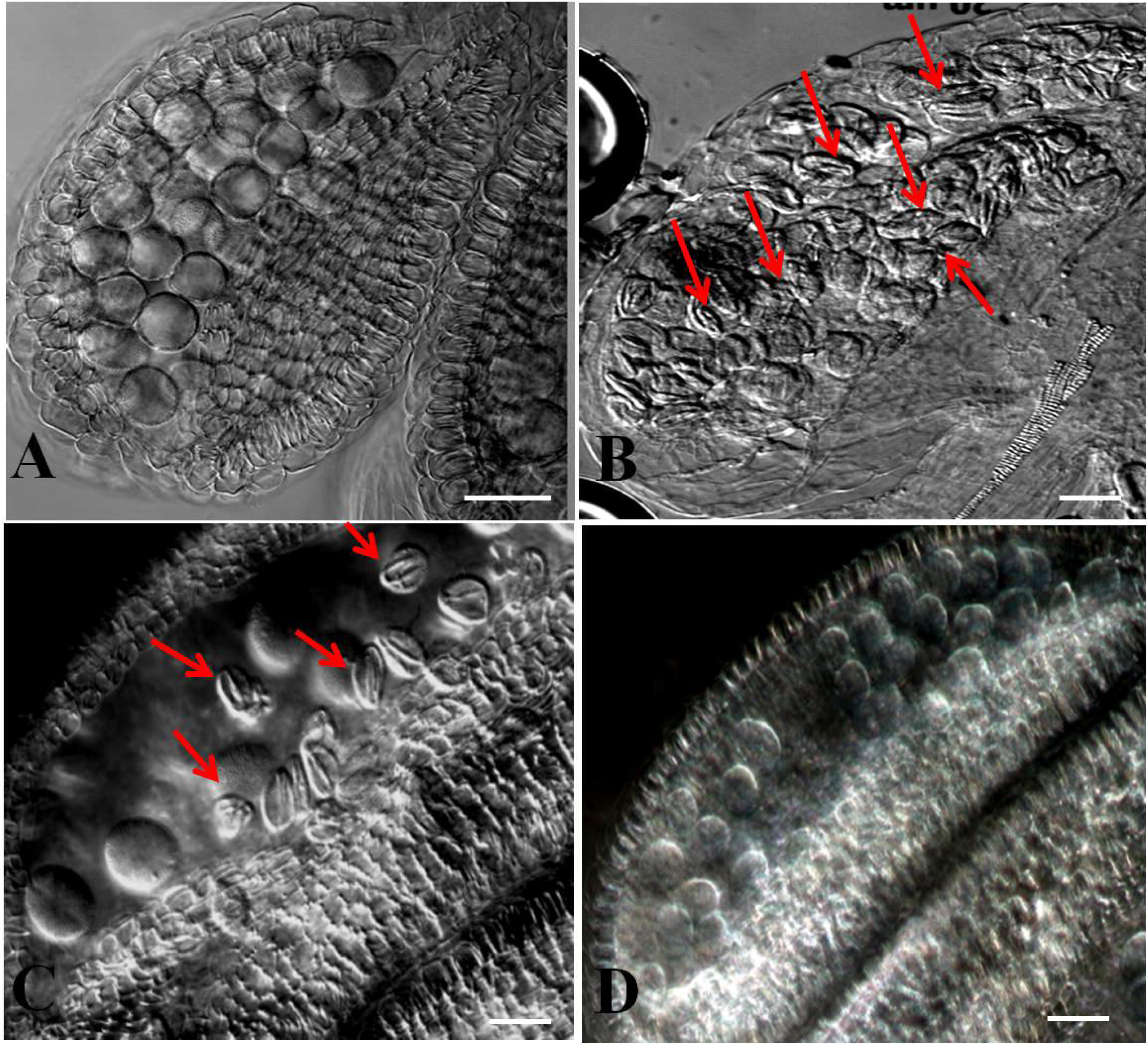
Phenotype of pollen in *Atnst1* mutant and complemented plants under DIC microscopy. (A) WT, (B) *Atnst1-1* mutant, (C) *Atnst1-2* mutant. Defective and collapsed pollen shown with arrows. (D) Anther of complemented plant filled with round pollen.

**Figure 7:**
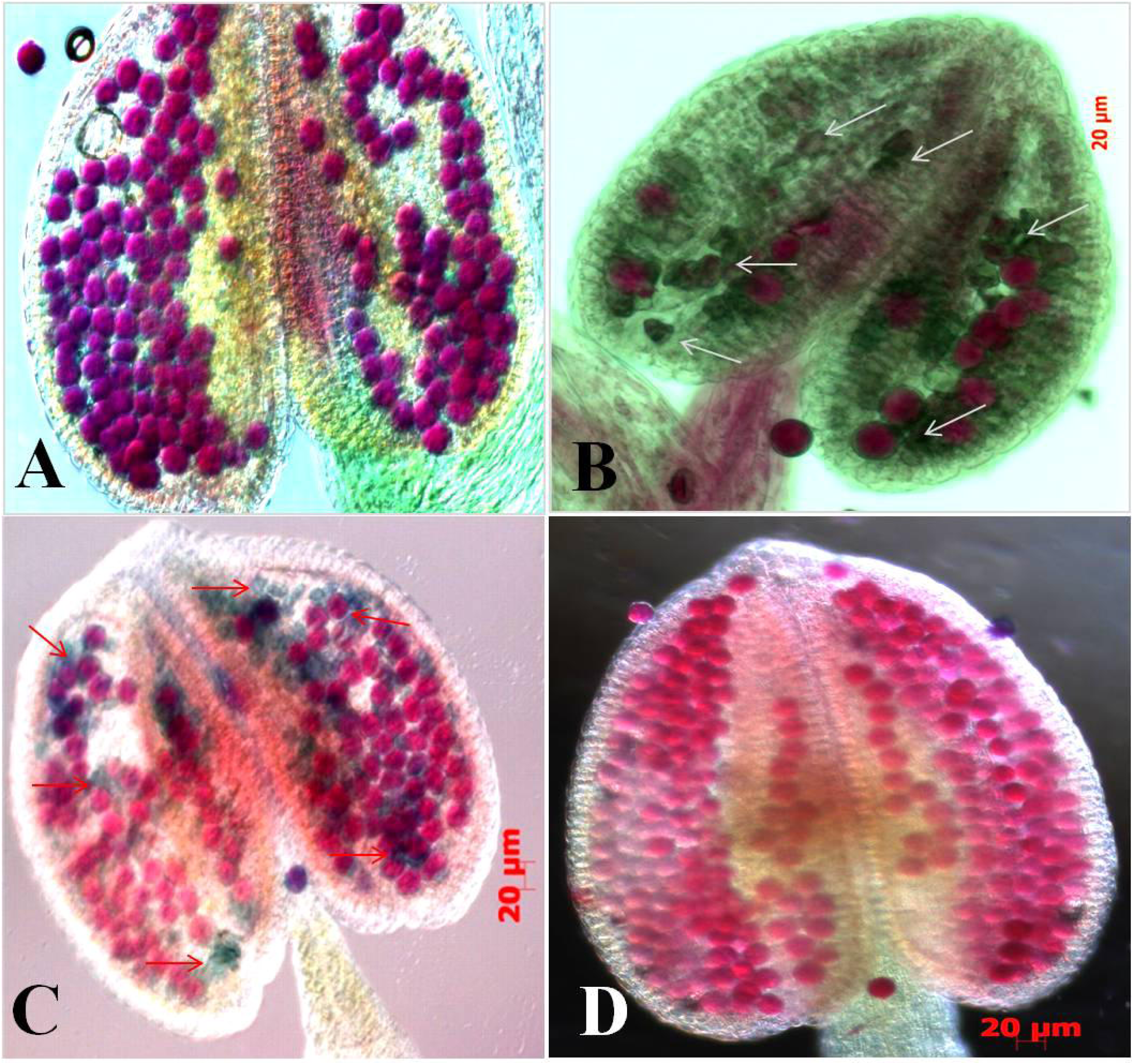
Assessment of pollen viability using Alexander’s staining. (A) WT, (B) *Atnst1-1* mutant, (C) *Atnst1-2* mutant, (D) complemented *Atnst1-1* mutant plant. Nonviable and collapsed pollen are indicated by arrows.

**Figure 8:**
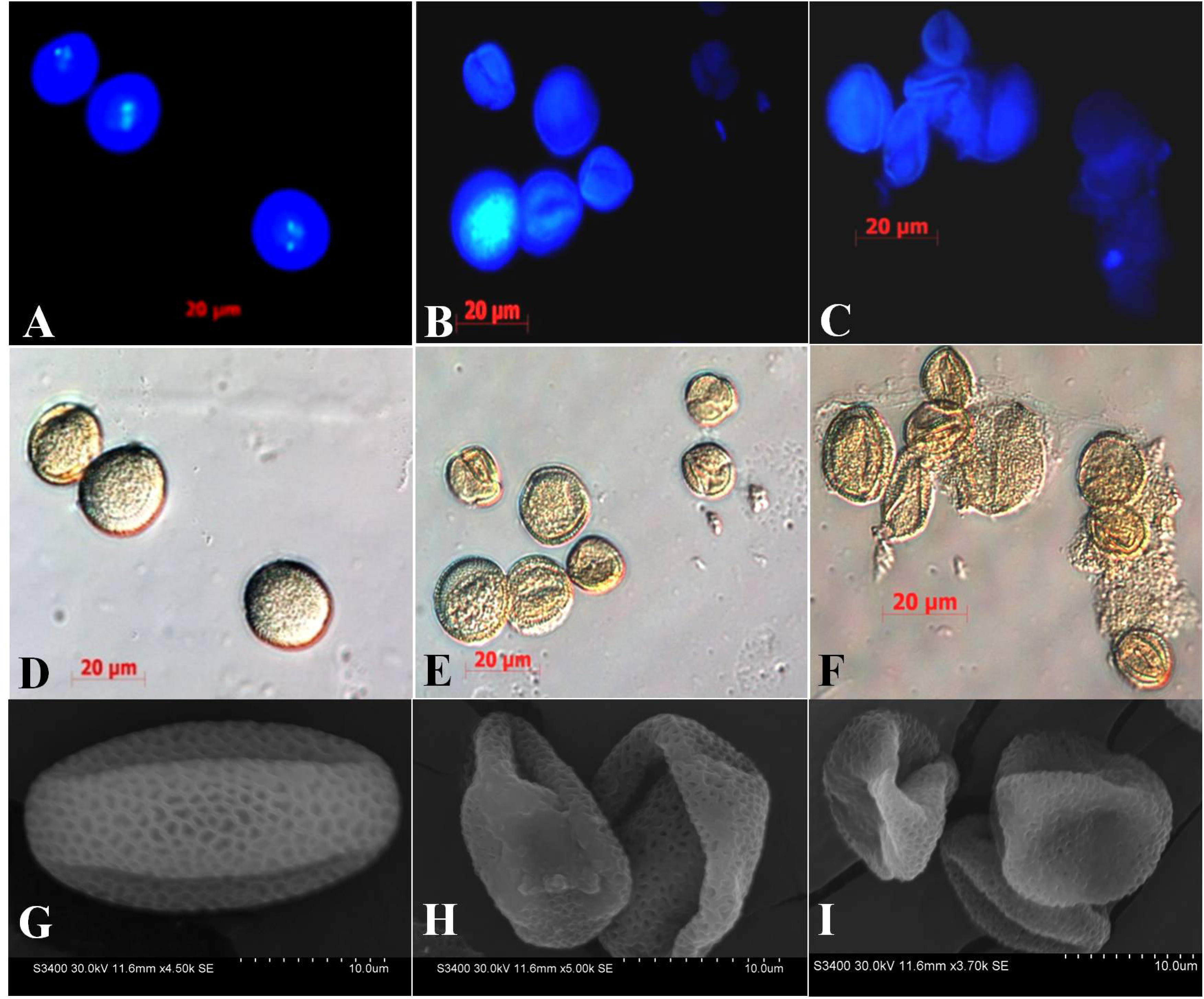
DAPI staining of mature pollen from (A) WT, (B) *Atnst1-1* and (C) *Atnst1-2* mutant plants. Collapsed pollen showed diffused or no nuclei, whereas wild type pollen contained normal tri-nuclear pollen. (D-F) DIC bright field images of the DAPI stained pollen. (G-I) Scanning electron micrographs of pollen from WT (G), *Atnst1-1* (H) and *Atnst1-2* (I) showing normal reticulate pattern of outer exine wall in normal as well as in collapsed pollen grains.

**Figure 9:**
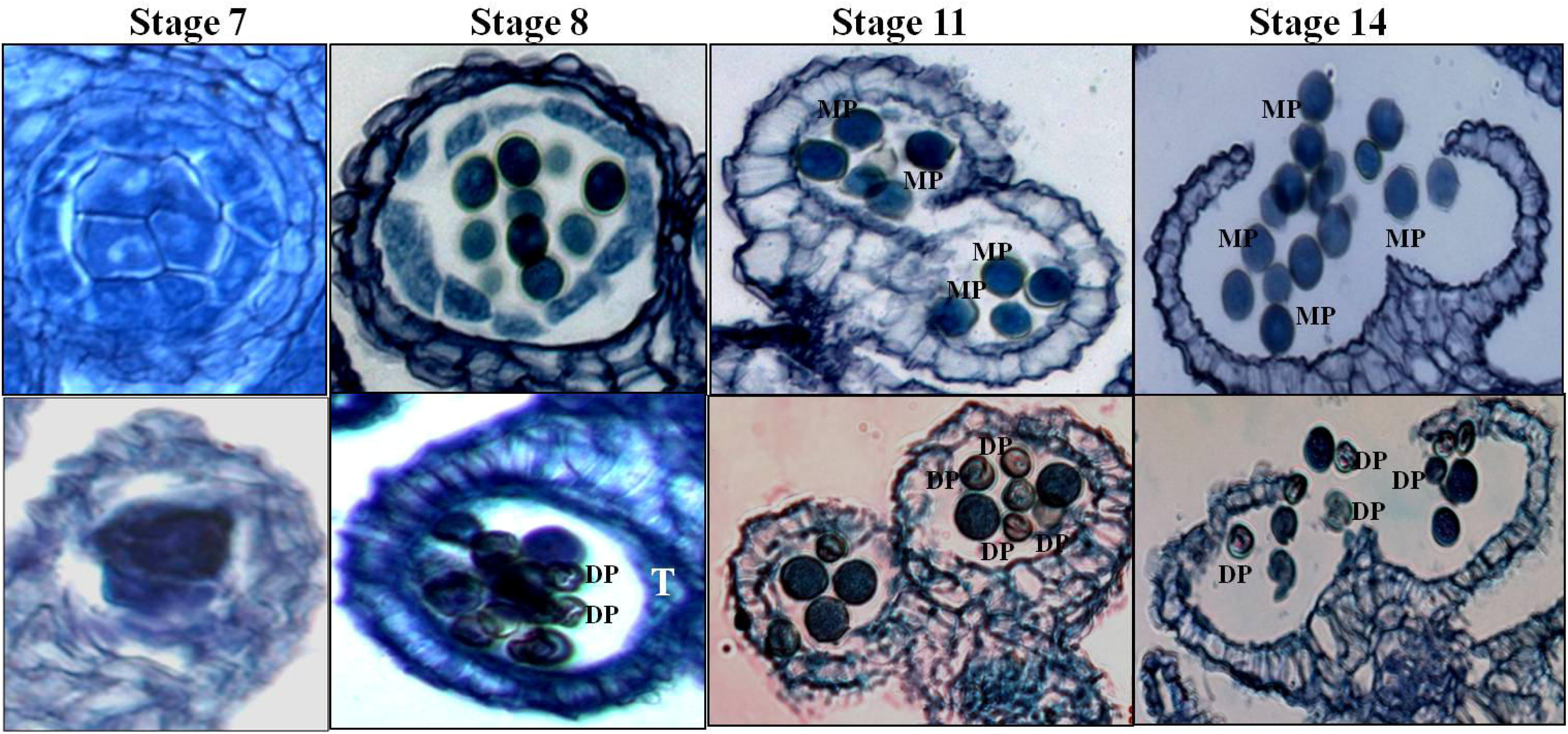
Histological study of pollen development in WT (upper panel) and *Atnst1* mutant (lower panel) *Arabidopsis* plants. Cross section of anthers at Stage 7 showing microspore mother cell (MMC). Stage 8 showing developing microspores and the tapetum in WT, whereas microspores with defective exine wall formation in *Atnst* mutant. Anthers at Stage 11 showing mature pollen and degenerated tapetum in WT, sterile and fertile pollen and partially consumed tapetum in mutants. Anthers at Stage 14 showing dehiscence and release of normal pollen in WT, whereas, the mutant shows defective and normal pollen. DP: defective pollen, MP: mature pollen, T: tapetum.

**Table 4.** Percentage of nonviable and aborted pollen in *Atnst* mutant.

| **Genotype** | **Total number of**<br>**pollen checked** | **Viable pollen (%)** | **Non- viable pollen (%)** |
| --- | --- | --- | --- |
| **WT** | 523 | 94 | 6 |
| *Atnst1-1*<br>(heterozygous) | 412 | 32 | 68 |
| *Atnst1-1* (homozygous) | 347 | 16 | 84 |
| *Atnst1-2*<br>(heterozygous) | 403 | 34 | 66 |
| *Atnst1-2* (homozygous) | 398 | 19 | 81 |

### Embryo sac development arrest at single nucleate stage in *Atnst* mutant

Results of reciprocal crossing confirmed that both male and female gametes were transmitting the mutant *At3g11320* allele but their inherent developmental defects lead to high frequency of failure of seed development. Therefore, female gametophyte development was also examined in the *Atnst* mutant line. Embryo sac development can be divided into two processes megasporogenesis and megagametogenesis. During megasporogenesis, the diploid megaspore mother cell undergoes meiosis and gives rise to four haploid nuclei.

In order to identify exact defects, stage-wise analysis of embryo sac development was documented with DIC microscopy of the tissue cleared ovules (Fig. 10). In mutant ovules, megaspore mother cell (MMC) differentiation, meiosis, degradation of three micropylar haploid megaspores and formation of functional megaspore appeared to be comparable to wild type (Fig. 10A and Fig. 10C). After this, 80% of the mutant ovules were found to arrest at the single nuclear stage and no further progress was observed in these ovules (Fig. 10D). This indicates that in these embryo sacs development is arrested at functional megaspore stage after differentiation (Fig. 10C). Functional megaspore remains as such and no further gametogenesis occurred. In remaining 20% of the ovules further gametogenesis progressed normally to give rise to seven-celled functional female gametophytes.

**Figure 10:**
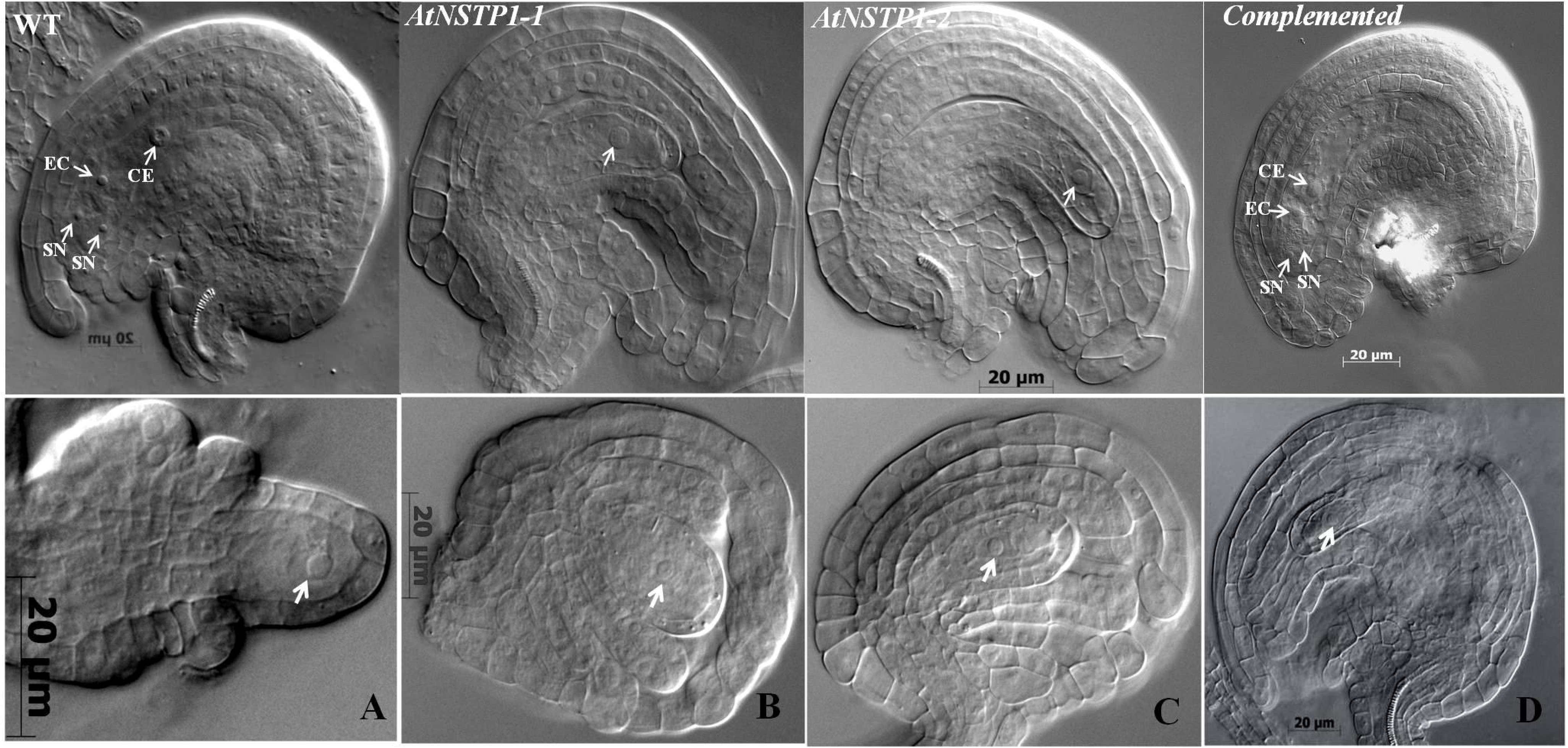
Ovule abnormalities of Arabidopsis *Atnst* mutants. WT ovule at anthesis showing fully differentiated embryo sac, *Atnst1-1* and *Atnst1-2* mutant ovules showing a single nucleate megagametophyte (arrows). Ovule from the plant complemented with the cDNA of *Atnst* showing fully differentiated embryo sac. A) Arrow indicating the MMC, B) PCD of three haploid spores at the micropylar end, C) differentiation of functional megaspore and D) absence of the progression of functional megaspore. CC: central cell, E: embryo, EC: egg cell, ES: endosperm, and SN: synergid cell.

### cDNA of the *AtNST* completely complements *Atnst* mutation

In order to rescue the gametophyte development defects in the mutant phenotype, cDNA (927 bp) of *At3g11320* (*AtNST*) gene was cloned into pCAMBIA 1302 expression vector under CaMV35S promoter as a GFP fusion protein and used to transform homozygous *Atnst1-1* mutant plants (Fig. 1D and Fig. 1H). Diagrammatic representation of the expression construct preparation is depicted in Fig. 11. In these mutant plants, expression of *AtNST* cDNA was confirmed by RT-PCR (Fig. 11B). Further, the *Atnst1-1* mutant plants were fertile and set seeds. Both the ovule as well as pollen development were normal with silique length and morphology comparable to the wild type. Seed sterility was completely rescued. Pollens were spherical and round in shape and regained their viability as evident from Alexander’s staining (Fig. 7D). Further, DIC analysis of the tissue cleared embryo sacs and pollens confirmed that both the gametophytes developed normally (Fig. 6D and Fig. 10). These results together demonstrated that mutation in *AtNST* gene is responsible for the sterility in this mutant.

**Figure 11:**
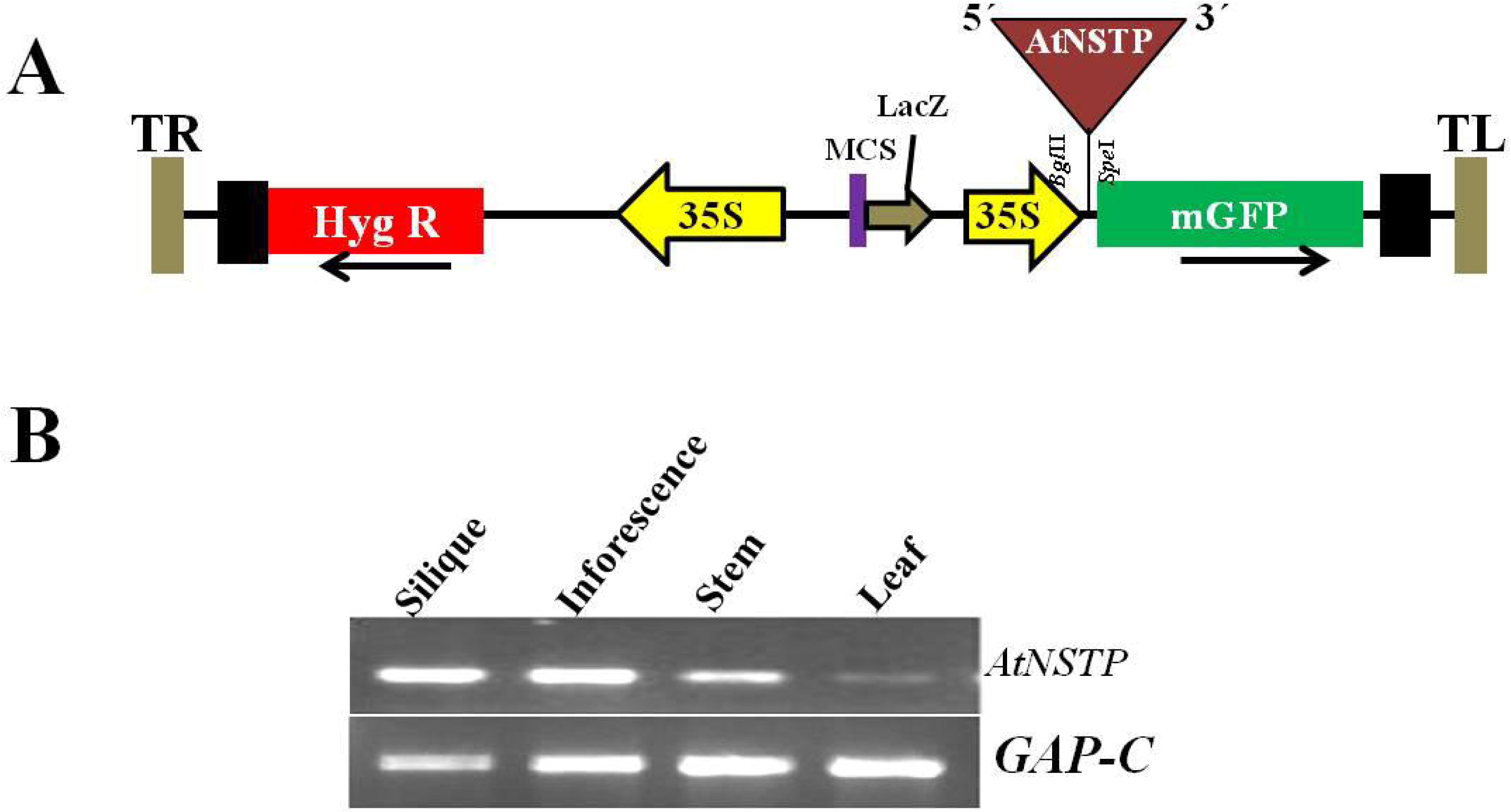
Complementation study of *Atnst1-1* mutant. (A) Diagrammatic representation of the construct prepared in pCAMBIA1302 for complementation studies. (B) RT-PCR analysis of *AtNST1* gene expression in complemented plants.

## Discussion

Glycosylation is one of the most common post-translational modifications of proteins with profound biological implications. NSTs which mediate the transport of sugars for this reaction play a central role in the glycosylation process. NSTs are the multiple domain, membrane spanning proteins which are present in the Golgi /ER membranes and help in translocation of nucleotide sugars from the cytosol to the lumen of the Golgi and endoplasmic reticulum (Handford *et al.,* 2006). During the glycosylation of proteins by the glycosyltransferases in the Golgi, these nucleotide sugars serve as a major source of sugar moieties. After glycosylation, nucleoside diphosphates are liberated and subsequently converted to nucleotides and are thought to exit the Golgi lumen through the action of nucleotide sugar-exchange transporters (Reyes & Orellana, 2008). According to Parsons *et al.,* (2013) the actual proteins responsible for these reactions have remained unknown. Some of the NSTs are highly specific in their substrate specificity and others can utilize three different nucleotide sugars containing the same nucleotide (Handford *et al.,* 2006). NSTs are not functionally redundant and play specific roles in glycosylation (Handford *et al.,* 2006). Reports are available that mutations in these genes cause developmental defects in Drosophila, *C. elegans* (Berninsone *et al*., 2001) and also alter the infectivity of *Leishmania donovani*. In humans, the mutation in GDP-fucose transporter causes defects in immune response and retarded growth. Role of Golgi nucleotide sugar transporter in modulation of cell wall biosynthesis in rice was reported by Zhang *et al.,* (2011). Niewiadomski *et al.,* (2005) reported the importance of glucose phosphate translocator (GPT) in pollen maturation and embryo sac development *via* oxidative pentose phosphate cycle mediated fatty acid biosynthesis in *Arabidopsis*.

In the present study, we identified a seed sterile mutant of *Arabidopsis* in which the mutation was in the *At3g11320* gene. T-DNA insertional mutation in this line involved unique transposition and rearrangement. The T-DNA first inserted in the *At3g11320* gene and then excised along with a part of 4^th^ exon of *At3g11320* gene, 3’ UTR and some part of intergenic region and reinserted inversely in the 2^nd^ intron of the *AtCYP90D1* (*At3g13730*) gene. Similar translocations and paracentric inversions following T-DNA insertions have been reported in earlier studies (Forsbach *et al.,* 2003; Lafleuriel *et al.,* 2004; Yuen *et al.,* 2005; Curtis *et al.,* 2009). It was expected that either of two genes (*AtCYP90D1*; *At3g13730* and *AtNST*; *At3g11320*) or both the genes may be responsible for the seed sterility phenotype. Analysis of independent SALK mutant lines having insertions at different positions of the respective genes resolved this and confirmed that deletion in *AtNST* gene is responsible for the mutant phenotype whereas T-DNA insertion in the *AtCYP90D1* gene did not show any clear phenotype.

*At3g11320* gene encodes a nucleotide sugar transporter family protein and shares homology with other transporter genes of *Arabidopsis* and rice at amino acid level. This protein contains 308 amino acids having a mass of 33.8 kDa with ten transmembrane domains. In the database this protein has been classified as triose phosphate translocator (TPT). TPT sugars can be used either for starch biosynthesis or for carbon skeleton of fatty acid biosynthesis. *In silico* analysis confirmed the presence of TPT domain in its sequence. Parsons *et al.,* (2012) characterized this protein as a Golgi membrane localized protein in a study of Golgi proteomics. Most of the known TPTs show subcellular localization in chloroplast. But despite the presence of TPT domain the AtNST protein encoded by *At3g11320* gene is localized in the Golgi membranes. Significant similarity of amino acids at the TPT domains in the *At3g11320* encoded AtNST might have led its grouping in the TPT/NST sub family.

RT-PCR data confirmed that *At3g11320* gene expression in almost all plant parts of the plant body including vegetative and reproductive parts in wild type *Arabidopsis*. But the mutation does not appear to affect vegetative growth, whereas the defects were observed in the male and female gametophytes only. In the *Arabidopsis* genome ∼40 *NST* genes are found and they share significant similarity among them. There may be subfunctionalisation among these NSTs and might account for specific defects in male and female gametophytes only. Alternatively, the demand of specific sugar moiety could vary among different tissues thereby leading to defects in specific tissues.

The data on reciprocal crossings confirmed that mutation of this gene affects both male and female gametes. Documentation of male and female gametophyte development in *Atnst* mutant showed collapsed, nonviable pollen (82%) at the maturity and single nucleate embryo sacs (80%) in the female gametophyte. Further, stage wise analysis of mega gametophyte development showed that meiosis of MMC is normal, after functional megaspore (FM) differentiation; it does not undergo mitosis and remains at single nucleate stage (Fig. 10). Similarly in male gametophyte also after releasing the pollen tetrads, they were unable to undergo further development and started degeneration ultimately they collapse (Fig. 9). Although the defects became evident in post-meiotic cells, actual functional impairment appears to set in earlier because more than 50% of the gametes in the heterozygous plant displayed defects in development.

In *Arabidopsis*, several starch-free mutants were characterised in which they retain the fertility and show no degenerated pollen grains (Kofler *et al.,* 2000). In plants, during gametophyte development there is a high demand of fatty acids (McConn *et al.,* 1996). During pollen grain development, after first mitosis accumulation of oil bodies and ER proliferation has been reported (Piffanelli *et al.,* 1998). In the present *Atnst* mutant both male and female gametophytes may be starved of triose phosphate sugars resulting in reduced supply of fatty acid precursors which affects lipid body formation and probably leads to impaired pollen wall formation and gametogenesis. Further, Saitoh *et al.,* (1996) reported the role of fatty acids in cell division. In yeast, they reported that *cut6* and *lsdl* mutants showed defects in nuclear division because of impaired fatty acid biosynthesis. Role of long chain fatty acids in endomembrane dynamics during cytokinesis was also reported in *Arabidopsis* (Bach *et al.,* 2011).

Several sugar transporters are expressed at different stages of pollen development (Truernit *et al.,* 1999; Schneidereit *et al.,* 2003; Scholz-Starke *et al.,* 2003; Schneidereit *et al.,* 2005). It was reported by Truernit *et al.,* (1999) that AtSTP2 transporter expression is confined to the early stages of pollen development i.e. at the beginning of callose degradation and microspore release from the tetrads. *AtUTR1* and *AtUTR2* mediated uptake of UDP-glucose into the ER was reported in *Arabidopsis* by Reyes *et al.,* (2010). Knocking out of both genes leads to abnormalities in both male and female germ line development. They also confirmed that these nucleotide sugar transporters are essential for the incorporation of UDP-glucose into the ER for the pollen development and embryo sac progress. According to the study of Parson *et al.,* (2012) this AtNST protein expression was localised as a Golgi membrane transporter protein. With this evidence we hypothesise that this nucleotide sugar transporter might be functioning to transport TPT sugars from cytosol into the Golgi lumen. These sugars might be required for the glycosylation of proteins/carbohydrates/lipids which might be required for the proper progression of pollen and embryo sac development in *Arabidopsis*.

The mutant phenotype including silique size and seed set was completely rescued when the cDNA of *AtNST* gene was over-expressed in the mutant back ground. Only a few (5-6%) seeds present at the terminal part of the siliques were found brownish and shrunken. Their phenotype was quite different from mutant phenotype. No abortive ovules were found in the siliques from the complemented plants. We conclude that the *AtNST* (*At3g11320*) gene is responsible for maintaining and regulating supply of sugars for the glycosylation of proteins/carbohydrates/lipids etc which are necessary for both male and female gametophyte development and thus plays an important role during plant reproduction in *Arabidopsis*.

## Acknowledgements

The authors thank the Indian Council of Agricultural Research, New Delhi, India, for financial assistance through the National Agricultural Innovation Project (NAIP/C4/C1072/2006-07) and are grateful to the Council of Scientific and Industrial Research (CSIR), New Delhi, India, for providing infrastructural and financial support in the form of projects MLP-072 and BSC-0107. RD acknowledges CSIR, New Delhi for the Junior and Senior Research Fellowships.

## Conflict of Interest

The authors have no conflict of interest

## Author Contribution statement

Rimpy Diman and Pininti Malathi : Performed experiments

Ramamurthy Srinivasan : Data analysis and manuscript preparation.

Shripad Ramchandra Bhat : Data analysis and manuscript preparation.

Yelam Sreenivasulu : Developed the concepts and approaches, performed experiments, data analysis and prepared the manuscript

## Supplementary figure legends

Figure S1: Identification of T-DNA insertion in *Atnst* mutant by Genome Walking. A. Diagrammatic representation of T-DNA insertion and locations of primers used for the confirmation of T-DNA insertion. B. Amplification of different sized of amplicons in the primary and nested PCR reactions in Genome Walking.

Figure S2: Diagrammatic representation of T-DNA insertion at the genome level after sequencing of amplicon and subsequent BLAST search against Arabidopsis genome and confirmation of insertion location by PCR with the primers designed corresponding T-DNA and to plant sequences flanking the T-DNA.

Figure S3: Diagrammatic representation of probable T-DNA translocations in Bitrap-636 mutant.

Figure S4: Diagrammatic representation of T-DNA insertion locations in *AtNST1-1*and *AtNST1-2* mutants, and B) confirmation of T-DNA insertions in the mutant lines through PCR.

Figure S5: Expression pattern of *AtNST* gene in WT Arabidopsis plants as shown in eGFP and GENEVESTIGATOR browsers.

Figure S6: Cellular localization of *AtNST* expression in onion epidermal cells as a function of GFP expression.

## Notes

### Competing Interest Statement

The authors have declared no competing interest.

