## Supplementary figures and images for "Loss of Nucleotide Sugar Transporter (*AtNST*) gene function in the Golgi membranes impairs pollen development and embryo sac progression in *Arabidopsis thaliana*"

### SF1

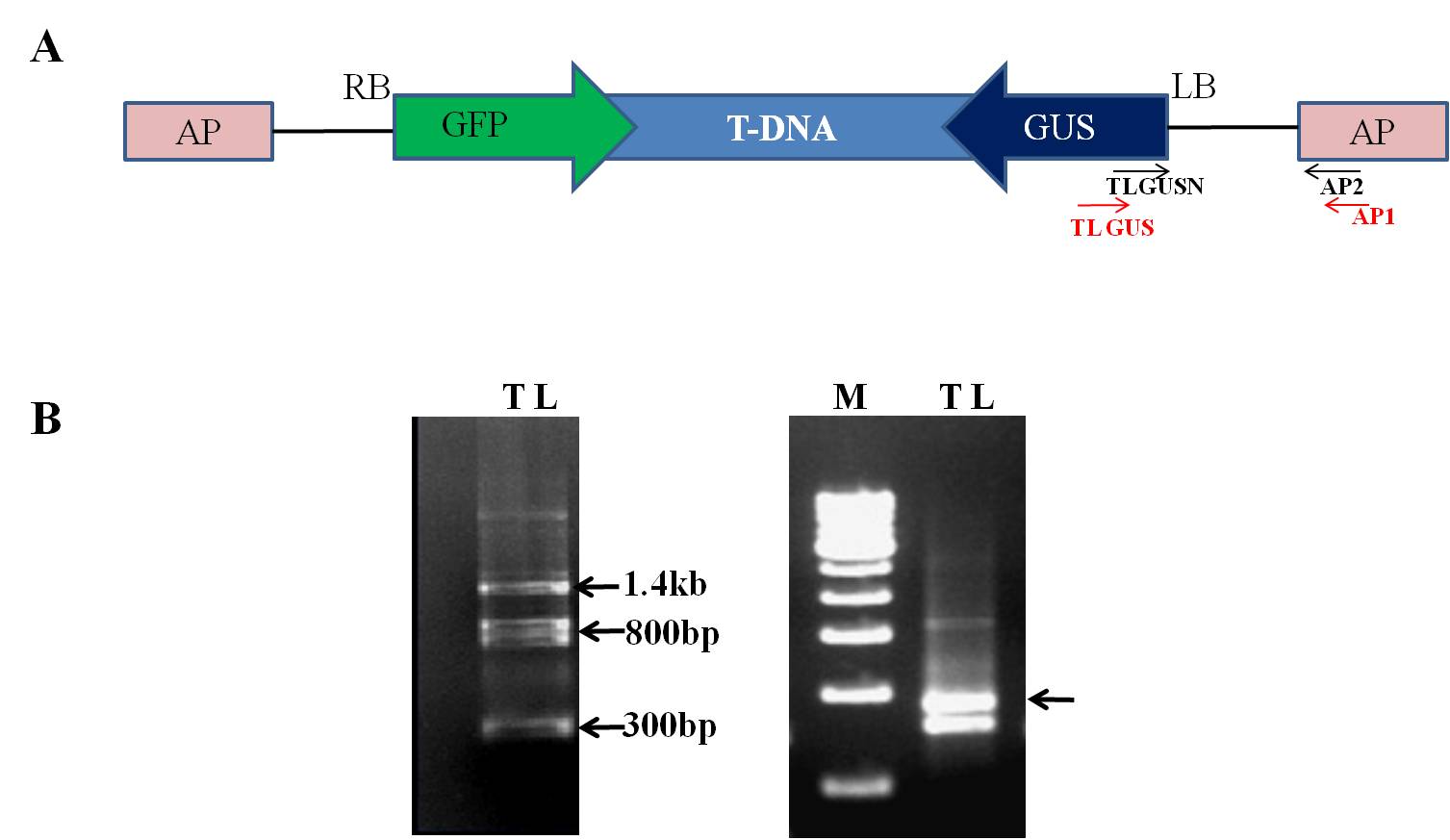

### SF2

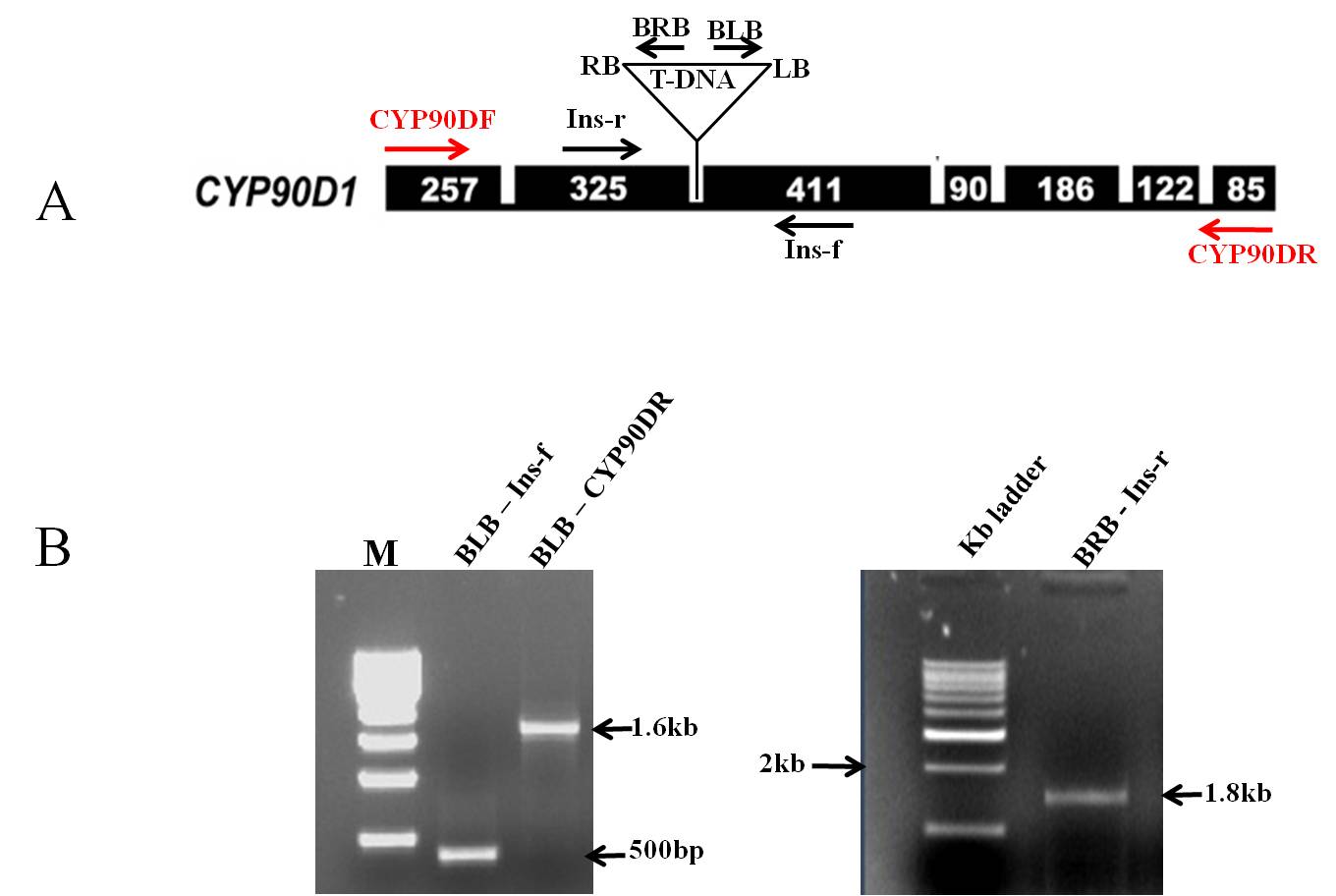

### SF3

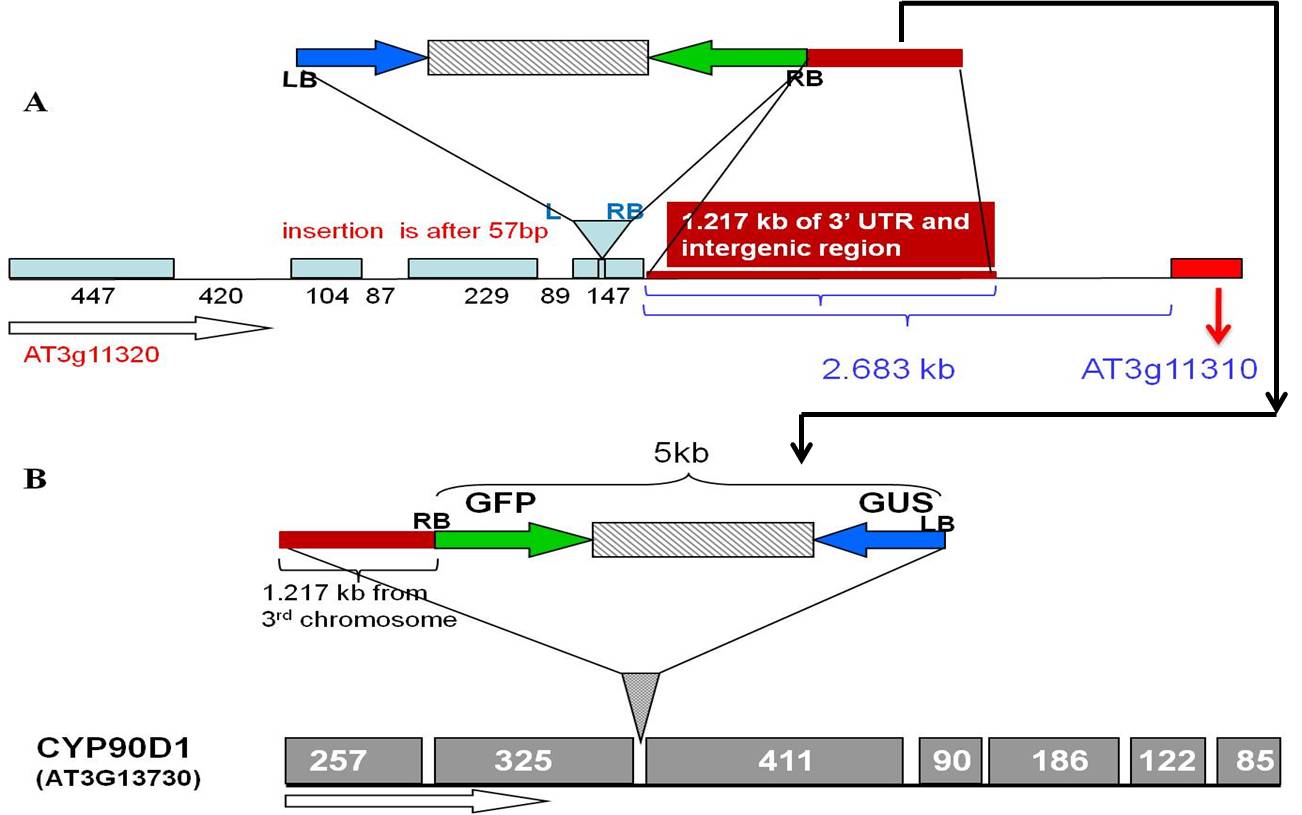

### SF4

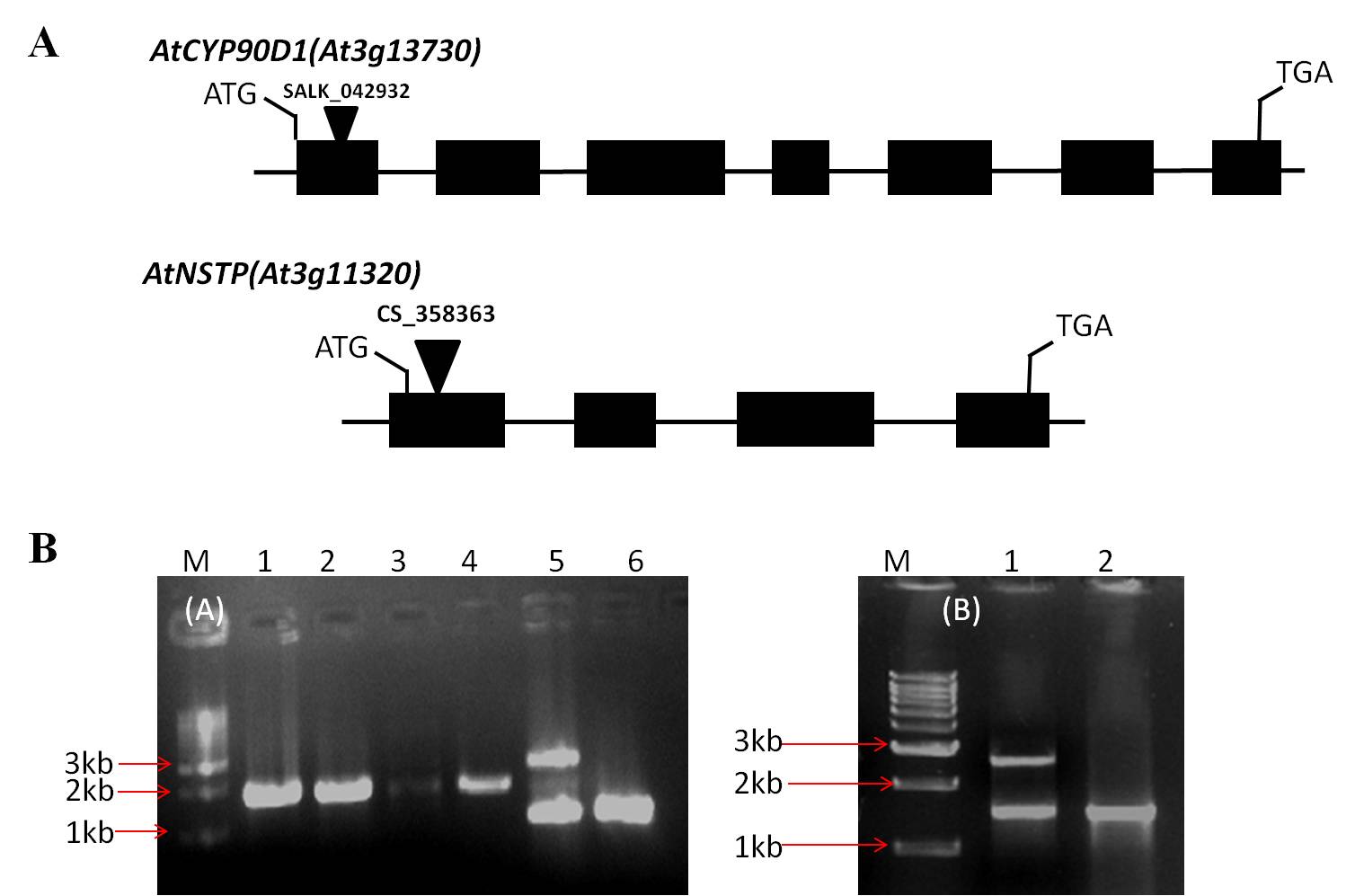

### SF5

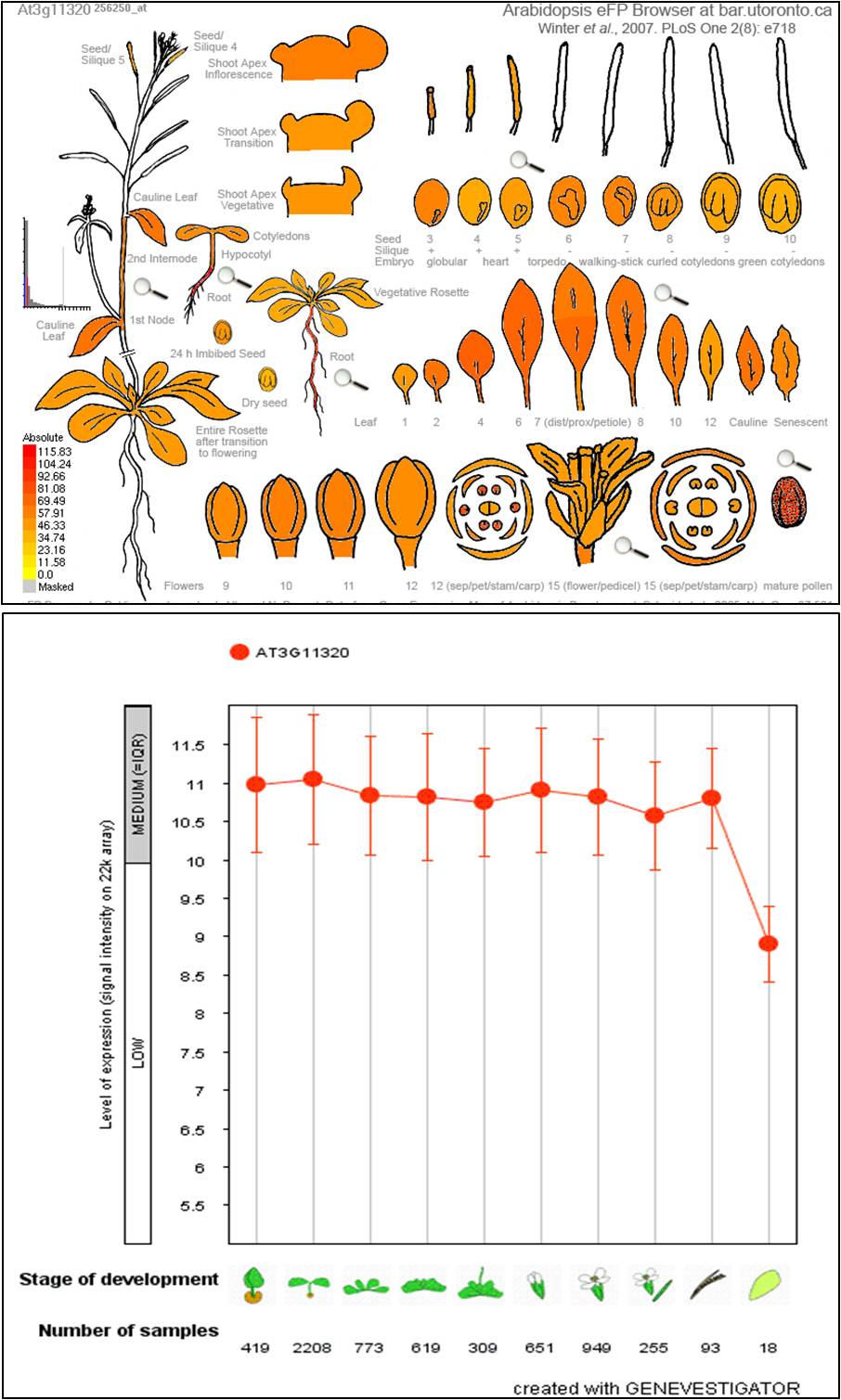

### SF6

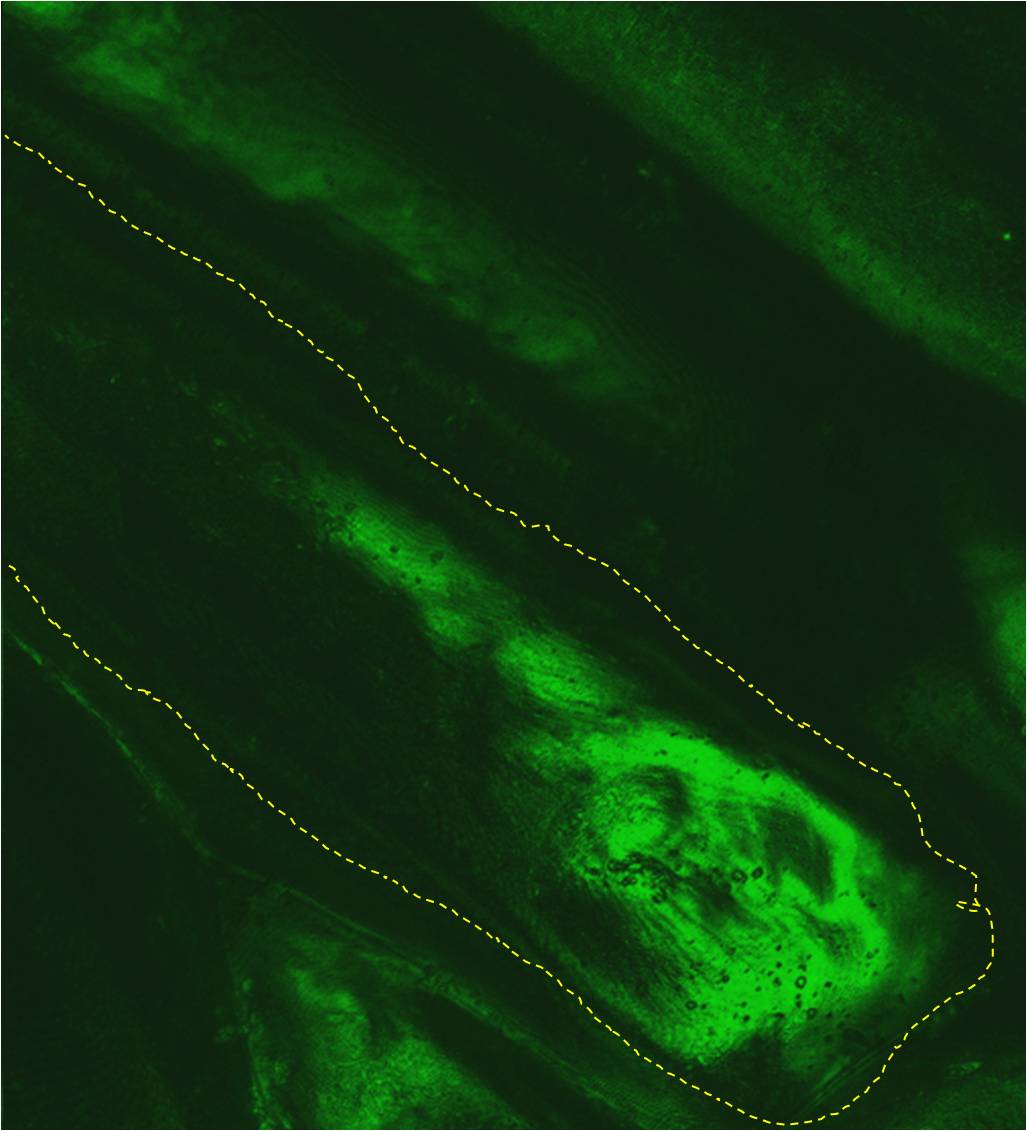
